# Intranasal administration of isradipine preferentially targets the brain

**DOI:** 10.1101/2025.03.10.639346

**Authors:** Jaime N. Guzman, Ema IIijic, Jack Nguyen, D. James Surmeier

**Affiliations:** Department of Neuroscience, Feinberg School of Medicine, Northwestern University, Chicago, Illinois 60611, USA; Cavalon Therapeutics, 2436 Oregon Street, Berkeley, CA 94705, USA

## Abstract

Parkinson’s disease (PD) is the second most common neurodegenerative disease. Despite a concerted effort on the part of the scientific community, there is no proven strategy for slowing PD progression. Nevertheless, there are several potential drug targets that if functionally modified could alter disease course. Preclinical, epidemiological and clinical trial data suggest that Ca_v_1 Ca^2+^ channels are one such target. Dihydropyridines (DHPs) are voltage-dependent, negative allosteric modulators of Ca_v_1 Ca^2+^ channels that are approved for human use. However, the brain concentration of DHPs that can be safely achieved in humans with oral dosing is limited because of the widespread distribution of these channels, particularly in the vasculature. Intranasal administration of DHPs is a potential alternative delivery strategy that has been used with compounds that have similar limitations. To test the viability of this drug administration strategy, mice were intranasally or orally administered the DHP isradipine mixed in one of three vehicles. Plasma and brain concentrations of isradipine were then determined using liquid chromatography/mass spectroscopy at subsequent times. These studies demonstrated that intranasal administration of isradipine was able to achieve higher brain concentrations than those in the plasma, and these differences persisted for hours. Thus, intranasal administration of DHPs could be used to achieve high levels of Ca_v_1 Ca^2+^ channel inhibition in the brain without producing unwanted peripheral side-effects.

## Introduction

Parkinson’s disease (PD) is the fastest growing neurodegenerative disorder in the world ^1^, compromising the quality of life and longevity ^2^. Although the cardinal motor symptoms of PD are effectively alleviated by levodopa therapy in the early stages of the disease, there is no proven strategy for slowing disease progression. One hypothesis about PD pathogenesis is that the loss of at-risk neurons is driven by sustained oxidant stress arising from the reliance upon feed-forward control of mitochondrial oxidative phosphorylation ^3^. Key nodes in this control network are plasma membrane, voltage-dependent Ca^2+^ channels with a Ca_v_1 pore-forming subunit ^4^. Dihydropyridines (DHPs) are negative allosteric modulators of these channels that have been shown to lower mitochondrial oxidant stress in at-risk neurons and to lessen the sensitivity to toxins and genetic mutations associated with PD. Moreover, epidemiological studies have consistently found that use of DHPs is associated with a reduced risk of developing PD ^5,6,7,8,9^. These observations led to a Phase 2 clinical trial that found at the maximum tolerated dose of a controlled release format of the DHP isradipine (10 mg/day), disease progression measured through clinical evaluation significantly slowed ^10^. However, a Phase 3 study using a similar daily dose of an immediate release (IR) format of isradipine failed to hit its clinical endpoints ^11^. That said, in patients that cleared IR isradipine more slowly, there were clear signs of disease slowing ^12^.

Taken together, these observations suggest that if the engagement of brain Cav1 Ca2+ channels by DHPs could be elevated and sustained, PD progression would slow. A major obstacle to achieving this goal is that DHPs are typically administered orally, leading to dosing limitations imposed by their action on the vasculature ^13^. An alternative route of administration that has been used with a number of drugs facing similar obstacles is through the intranasal mucosa ^14^. This route gives drugs with appropriate properties direct access to the brain, bypassing the gut and circulatory system. The studies described here were intended to determine whether isradipine falls into this category. Our studies revealed that if delivered with an appropriate vehicle, intranasal administration of isradipine achieved a high brain/plasma ratio that persisted for hours, pointing to a strategy for overcoming the limitations imposed by oral dosing.

## Methods

All animal care and procedures were performed according with Northwestern University Animal Studies committee and in accordance with the National Institutes of Health Guide for the Care and Use of Laboratory Animals. Male and female C57BL/6 mice (3 months old and ∼ 25 g) were purchased from Jackson Laboratory (Bar Harbor, Maine). Isradipine suspensions with final concentration of 25 mg/ml (67.4 mM) at 20-25°C were prepared in vehicle 1 (0.5% carboxymethylcellulose (CMC) in saline) or in vehicle 2 (N,N-dimethylformamide, PEG 400, in saline (2:6:2 ratio)). Isradipine also was prepared in dimethylsulfoxide (DMSO) vehicle at concentration of 50 mg/ml (134.6 mM) at 20-25°C.

Under light isoflurane anesthesia, mice received either 2-3 intranasal doses or one oral dose of isradipine to achieve a final dose of 5 or 10 mg/kg (administered volume was different for each animal based on their weight). See attached video. Animals were returned to the cage and then sacrificed at different time points. At these time points, mice were terminally anesthetized with a mixture of ketamine (50mg/kg)/xylazine (4.5mg/kg). For determination of isradipine in the plasma, 400-50 µl ml of blood was drawn from anesthetized mice from posterior vena cava and incubated at room temperature in EDTA-coated vials for 30 min. After centrifugation, supernatants were collected, transferred to capped polypropylene test tubes, and stored at −80°C until the day of the assay. Brains were collected after decapitation and fast frozen on dry ice and stored at −80°C. All samples were wrapped in aluminum foil during the procedure.

There were three different conditions for which the isradipine concentration was determined. First, brain and plasma isradipine concentration were determined 30 minutes after intranasal drug administration. These assays compared vehicle 1 and 2 at two doses (5 and 10 mg/kg). Mice in this trial (N=15) were given 2-3 µl of the isradipine suspension per nare. As isradipine was not completely soluble at the concentrations used, the suspension was well mixed before dosing. Second, to determine brain and plasma isradipine concentration at different time points (0.5, 1, 3, 6 and 24 hours after administration), isradipine was administered in vehicle 2 at volumes that yielded a final dose of 15mg/kg. Mice (N=16) were sacrificed as described at the predetermined time points as described above. Lastly, the brain and plasma isradipine concentrations achieved using vehicle 2 were compared to those achieved with a DMSO vehicle. In these experiments, mice (N=48) were given 10 mg/kg isradipine intranasally and orally. Plasma and brains were collected 0.5, 1, 3 and 6 hours after dosing.

Data points at each time point and condition (n=3) were averaged and plotted.

## Results

In the first experiment, the impact of the vehicle used to administer isradipine was examined. Isradipine was given at two doses (5 and 10 mg/kg) and plasma and brain concentration were measured 30 min later. With vehicle 1 (0.5% CMC in saline), isradipine at a low dose (5 mg/kg) achieved a median brain concentration of 6 nM, whereas isradipine was not detected in the plasma; at a higher dose (10 mg/kg), brain isradipine concentration rose to a median of 36 nM, whereas it was 11 nM in plasma (Fig. 1A). With vehicle 2 (N,N-dimethylformamide, PEG 400, saline), isradipine at a low dose (5 mg/kg) achieved a median brain concentration of 244 nM, whereas the plasma median concentration was 55 nM; at a higher dose (10 mg/kg), the median brain concentration of isradipine rose to 732 nM, whereas the median concentration in the plasma was 332 nM (Fig. 1B). Thus, regardless of vehicle, intranasal administration of isradipine achieved higher brain than plasma concentrations. That said, vehicle 2 was clearly superior in delivery characteristics. Consequently, subsequent experiments used vehicle 2.

**Figure 1.**
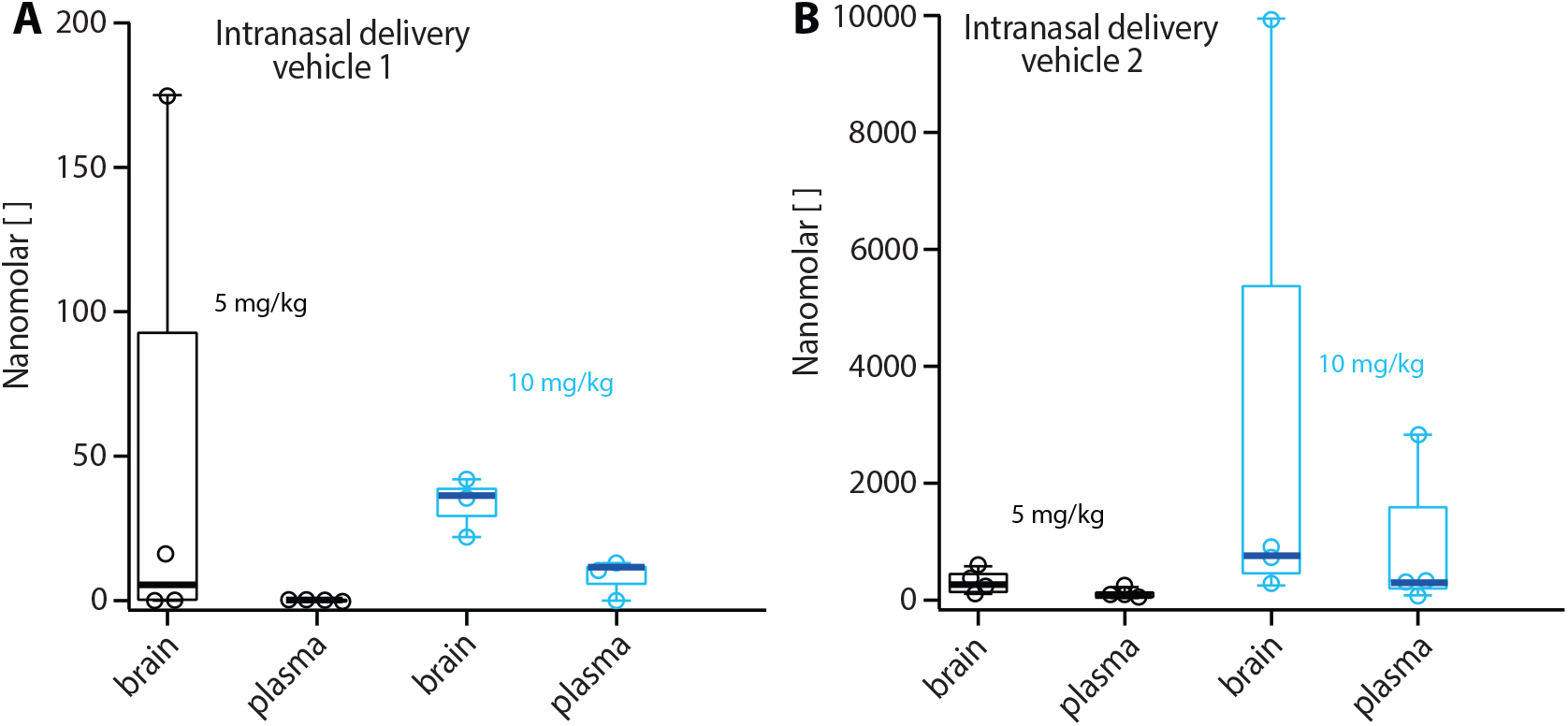
Brain concentrations were higher than plasma concentration after intranasal delivery of 5 and 10 mg/kg of isradipine in two different vehicles. (A) Box plots showing distribution of brain and plasma isradipine concentration in nanomolars 30 minutes after intranasal delivery of 5 mg/kg (black) and 10 mg/kg (blue) of isradipine suspended in 0.5% CMC in saline (vehicle 1). There were four mice in 5 mg/kg group and three mice in 10 mg/kg group. (B) Box plots showing distribution of brain and plasma isradipine concentration in nanomolars 30 minutes after intranasal delivery of 5 mg/kg (black) and 10 mg/kg (blue) of isradipine suspended in N,N-dimethylformamide, PEG 400, saline (vehicle 2). There were four mice per each group.

In the second experiment, brain and plasma isradipine concentrations were determined at regular intervals after intranasal administration at a dose of 15 mg/kg in vehicle 2. As shown in Figure 2, the isradipine concentration in the brain was higher than plasma at all time points. Interestingly, the brain/plasma concentration ratio rose from 2.5 at 30 minutes after administration, to 4 at 1 hour, and 80 at 3 hours. Isradipine was not detectable in the plasma 6 hrs after administration, while brain concentration was 30 nM. Twenty-four hours after administration, isradipine was not detectable in either the brain or plasma.

**Figure 2.**
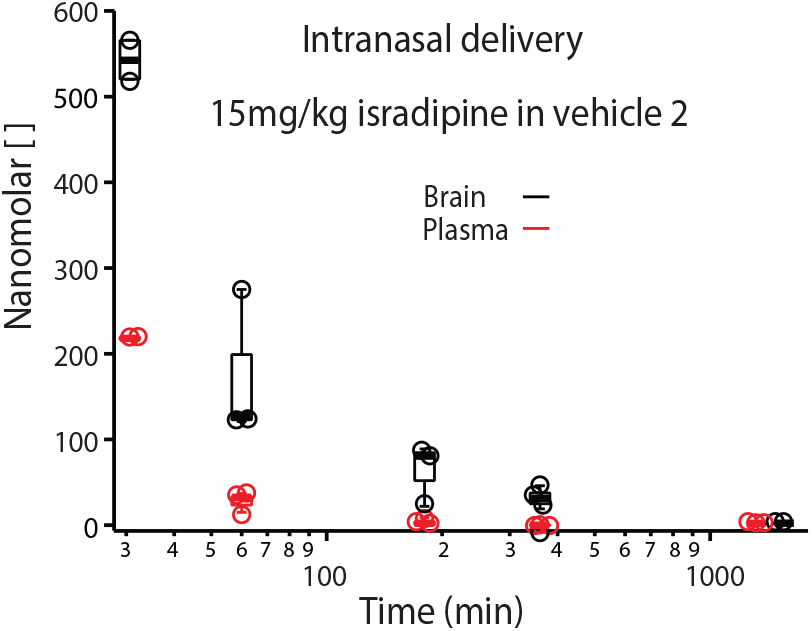
Brain concentrations were higher than plasma concentration after intranasal delivery of 15 mg/kg of isradipine in N,N-dimethylformamide, PEG 400, saline at all checked time points. Box plots showing distribution of brain (black) and plasma (red) isradipine concentration in nanomolars 30, 60, 180, 360 and 1440 minutes after isradipine administration. In the box plot showing isradipine concentration in brain after 30 min there were two mice samples because the third result was removed as a statistical outlier. All other box plots include data from three mice. Note that time is presented on logarithmic axis.

In the third experiment, intranasal isradipine administration was compared with oral administration (gavage). Isradipine (10 mg/kg) was administered either in vehicle 2 or in DMSO; in vehicle 2, isradipine was in suspension, whereas it was fully dissolved in DMSO. With DMSO as the vehicle, intranasal application of isradipine achieved brain concentrations that were 2-8 times higher than those achieved with the isradipine suspension in vehicle 2 (Fig. 3A, C). Oral administration of isradipine in DMSO achieved much lower brain concentrations (Fig. 3B). Unexpectedly, oral administration of isradipine in vehicle 2 achieved higher brain and plasma concentrations than intranasal delivery (Fig. 3D). However, even with this route of administration, brain concentrations of isradipine were higher than those in the plasma at all time points.

**Figure 3.**
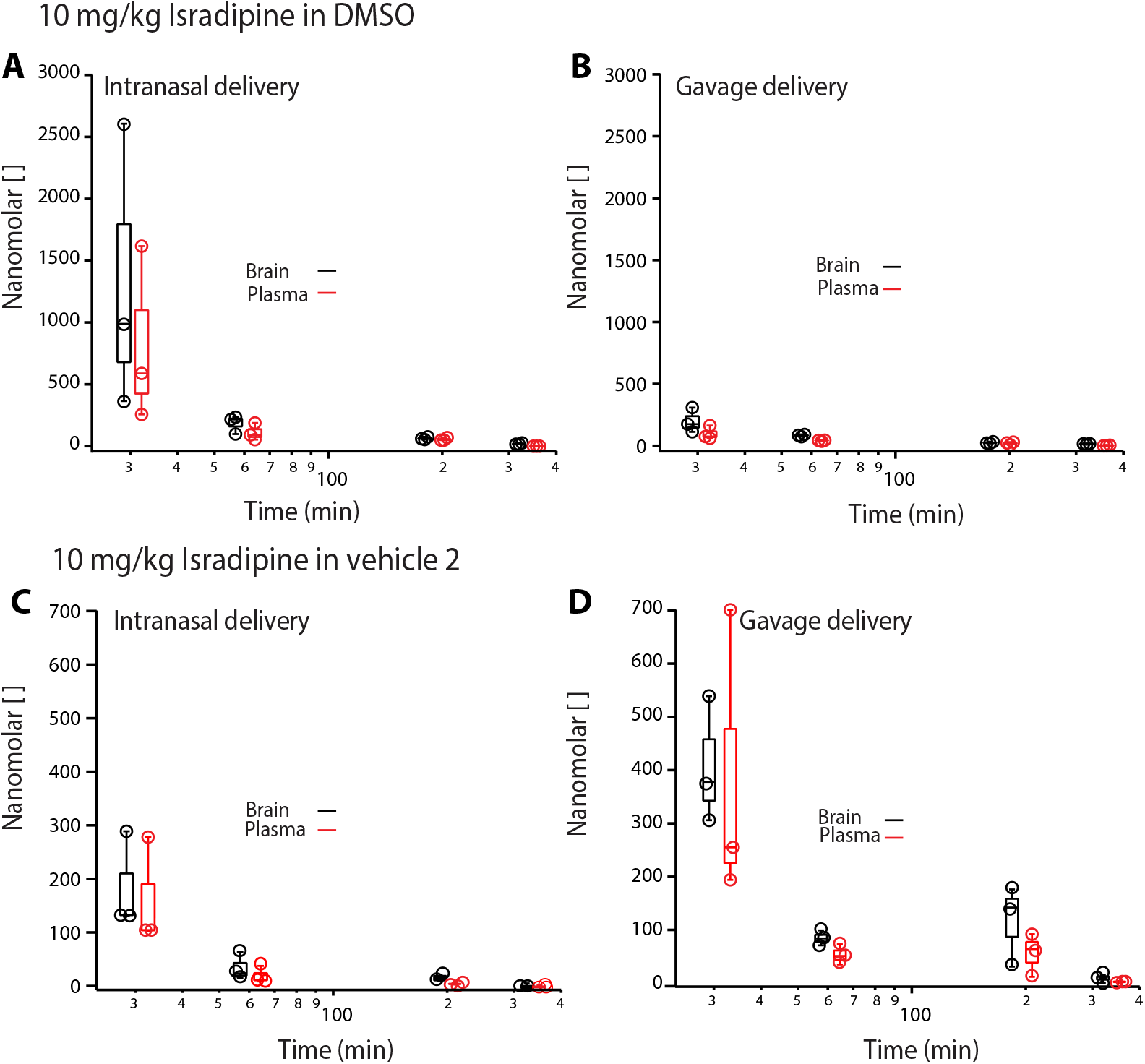
Brain concentrations were higher than plasma concentration after intranasal and oral delivery of 10 mg/kg of isradipine in DMSO and in N,N-dimethylformamide, PEG 400, saline at all checked time points. Box plots showing distribution of brain (black) and plasma (red) isradip-ine concentration in nanomolars 30, 60, 180, and 360 minutes after isradipine administration through intranasal (A) and oral (B) route. Isradipine was dissolved in DMSO and there were three mice in each group. Box plots showing distribution of brain (black) and plasma (red) isradipine concentration in nanomolars 30, 60, 180, and 360 minutes after isradipine administration through intranasal (C) and oral (D) route. Isradipine suspension was made in N,N-dimethylformamide, PEG 400, saline and there were three mice in each group except at 360 minutes after intranasal administration group where one data point was removed as a statistical outlier leaving two data point in that group. Note that time is presented on logarithmic axis.

## Discussion

The main finding of this study was that intranasal administration of the DHP isradipine was able to achieve higher brain concentrations than those in plasma at all time points examined (0.5-6 hrs), regardless of vehicle. That said, the vehicle used to deliver isradipine had a significant effect on the concentration of isradipine that was achieved. Three vehicles were tested. Both 0.5% CMC/saline (vehicle 1) and N,N-dimethylformamide/PEG400/saline (vehicle 2) were tested because of their use in other studies of intranasal drug delivery ^15,16^. At the concentrations necessary to achieve our dosing goals with 2-3 µl/nare, neither vehicle allowed isradipine to be fully dissolved and it was delivered in a well-mixed suspension. This limitation may account for some of the variability in the data. Whether this was a factor in the decidedly lower (∼10X) brain concentrations achieved with 0.5% CMC in saline compared to N,N-dimethylformamide, PEG400 in saline is unclear. Regardless, intranasal administration achieved brain concentrations of isradipine that were 2-3 times greater than those in the plasma.

Using an N,N-dimethylformamide/PEG400/saline vehicle, intranasal isradipine administration achieved a brain/plasma ratio that initially was around 3 and then rose to near 10 six hours later. Six hours after intranasal administration, the median brain concentration of isradipine was greater than 20 nM – within the range of concentrations previously shown to be protective against PD-related toxins ^17^. This pharmacokinetic profile likely reflects the portioning of amphipathic isradipine into the lipid phase of the brain.

To determine how getting isradipine completely into solution might affect transit to the brain, DMSO was used in a third set of experiments. Isradipine has a much higher solubility (∼50 mg/ml) in DMSO than either vehicle 1 or 2. With this combination, higher brain concentrations of isradipine were achieved than with vehicle 2; moreover, the brain/plasma ratio remained high. The extent to which this reflects the impact of isradipine solubility in DMSO or the ability of DMSO to promote the transit of isradipine into the brain is unclear. Answering this question and determining the tolerability of different solvents requires further study.

In the last series of experiments, intranasal administration of isradipine was compared to oral gavage. With DMSO as the vehicle for isradipine, oral gavage was much less effective in elevating brain concentrations – even though the brain/plasma ratio remained greater than 1. Unexpectedly, with the N,N-dimethylformamide/PEG400/saline vehicle oral gavage was more effective in elevating brain isradipine concentration than intranasal administration. It is unclear why this was the case. Interestingly, even in this situation, brain concentrations of isradipine were higher than those in the plasma.

Although much remains to be determined, our studies suggest that intranasal delivery of isradipine is a promising strategy for achieving neuroprotective concentrations of isradipine (and other DHPs) in the brain. Previous studies have shown that peripheral administration of DHPs dose-dependently protects dopaminergic neurons in toxin-based animal models of PD ^17–19^. Both dopaminergic cell bodies and terminals were spared in toxin models when plasma isradipine concentrations reached 10-20 nM ^17^. However, in this plasma concentration range, many humans report adverse events, including edema, dizziness and fatigue ^20^. In principle, intranasal administration of isradipine could be used to achieve brain concentrations in the protective range (10-20 nM), while keeping plasma concentrations below the threshold for significant side-effects. However, given that isradipine is eventually cleared from the brain, maintaining isradipine within a protective range would require repeated, periodic intranasal dosing. The advent of calibrated, easy-to-use intranasal delivery devices ^21^ could make this requirement relatively easy to surmount.

## Supporting information

See attached video

## Acknowledgments

This work was supported by a grant from the Cure Parkinson’s Trust and the JPB Foundation. We also wish to acknowledge Dr. Ben Owen, PhD, Director of the Clinical and Pharmacology Core, Department of Chemistry, Northwestern University.

